# Inferring the nature of missing heritability in human traits

**DOI:** 10.1101/373290

**Authors:** Eugenio López-Cortegano, Armando Caballero

## Abstract

Thousands of genes responsible for many diseases and other common traits in humans have been detected by Genome Wide Association Studies (GWAS) in the last decade. However, candidate causal variants found so far usually explain only a small fraction of the heritability estimated by family data, the rest remaining ‘missing’. The most common explanation for this observation is that the missing heritability corresponds to variants, either rare or common, with very small effect, which pass undetected due to a lack of statistical power. We carried out a meta-analysis using data from the NHGRI-EBI GWAS Catalog in order to explore the distribution of gene effects for a set of 59 complex traits, to investigate the factors related to new gene discovery and to quantify their contributions to heritability. With the data obtained we were able to predict the expected distribution of gene effects for 16 traits and diseases including cancer and immune disorders, the missing number of genes yet to be discovered, and the additive contribution of common variants to heritability. Our results indicate that, for most traits, the additive contribution of hundreds or thousands of genes is able to explain the familiar heritability. However, for some traits, the predicted heritability is lower than the familiar one, so that part of the missing heritability cannot be explained by the contribution of common variants with additive effects, and other phenomena (*e.g*. dominance or epistasis) should be invoked.

**AUTHOR SUMMARY:** The heritability of biological traits refers to the fraction of the phenotypic (*i.e.* visible or diagnosable) variability that is explained by the underlying genetic variability. Despite the efforts made to find the genes responsible for complex traits and diseases, most of the heritability explained by the variants found explains only a small fraction of that measured by population and family studies. The remaining ‘missing’ heritability is usually assumed to be due to the additive effect of undetected variants. Here we carried out a meta-analysis using records from a publicly available database of genome-wide studies on humans. With these data, we were able to investigate and make inferences on the nature of heritability and the factors associated to new gene discovery. Our results indicate that increasing population sample size, as well as its diversity, enhances the discovery of new genes, but these have lower and lower effects, contributing little to heritability. We were also able to predict the distribution of gene effects for many traits and the number of variants needed to fully explain the heritability. For some traits, the additive effects of single variants yet to be discovered cannot explain the heritability, suggesting that other sources of variation are involved.

## INTRODUCTION

Understanding the genetic architecture of complex traits has become a fundamental topic of study in human genetics(1). In the last years huge efforts have been made to investigate the genetic basis of human complex traits through Genome-Wide Association Studies (GWAS) or meta-analyses of their results(2–5). There has been a parallel increase in the number of big Consortiums able to carry out large GWAS with higher and higher numbers of individuals and, therefore, with increasing statistical power(6,7), as well as of genomic repositories, including databases specialized in published GWAS results(8). Up to date, more than 15,000 unique SNPs have been identified to be associated with human diseases or other traits with genomic significance, according to data recorded by the NHGRI-EBI GWAS Catalog(8). However, SNP markers of known variants cannot explain but a small percentage of the heritability measured by cohort studies for almost every studied trait, what has been referred to as ‘missing’ heritability(9).

The most common assumption to explain the missing heritability is that many common variants of small effect pass unnoticed in most GWAS due to a lack of statistical power(10), and a number of variants on the order of hundreds to thousands are yet to be found(5). In fact, the missing heritability gap of well-studied traits such as human height is reduced when GWAS are performed with increasing sample sizes (*i.e*. higher statistic power) (6). However, the newly found SNPs may have smaller effect sizes on average(11), so that their contribution to heritability may be small when compared to that of genes of moderate effect size. In addition to the additive effect of common or rare variants, many other genetic phenomena, such as epistasis(12), epigenetics(13) or dominance(14), among others, have been suggested to be involved in the explanation of missing heritability. Thus, prior to find the missing heritability, it is indispensable to better understand the genetic architecture of common traits and diseases.

Here we performed a meta-analysis using the GWAS Catalog(8) in order to explore the genetic basis of human complex traits, and to better understand the nature of missing heritability for a number of human diseases and common traits, focusing on: i) the detection of factors related to new gene discovery by GWAS, ii) investigating the nature of the distribution of gene effects sizes, and how these genes contribute to heritability, and iii) the prediction of expected additive contributions to heritability. Our results indicate that, while the heritability of some complex traits can be explained by the joint additive effect of hundreds to thousands variants, in agreement with previous works, for some other traits the missing heritability gap can only be filled by non-additive effects, suggesting a role of non-additive genetic variance in explaining the missing heritability.

## RESULTS

### Factors associated to gene discovery and nature of the variants

We first investigated which are the main factors associated with GWAS gene discovery for a set of 59 traits (Table S5 in Appendix 1 in Supplemental Information), concluding that population sample size is the main one (*P* < 2 × 10^−16^). Publication year was also significant, but with higher *P*, and was significantly correlated to population size (*r* = 0.15, *P* = 1.36 × 10^−3^) as expected, as more recent studies usually have larger sample sizes. The regression of the number of genes found on sample size is rather variable between traits (Figure S1 and Table S5 in Appendix 1), but on average, suggest that a sample on the order of 53,000 individuals is required to start detecting genes, and that 6,000 individuals are needed to incorporate each new gene, though these numbers range widely across traits. Interestingly, we also find an association of the number of genes discovered with the number of continents of origin considered alone (*b_ncont_* = 3.37; *p* = 0.02), which is uncorrelated with population sample size (*r* = 0.08; *p* = 0.11).

Figure 1 shows that the average gene effect (α) steadily declines as new genes are found with larger samples sizes. The total number of genes analysed for each of the traits is shown in Table S2. The decline is remarkably consistent across traits, with a two-parameter exponential model fitting best the observations (average *R*^2^ = 0.96; see Table S3[A]). The trend observed is in accordance with the expectation that genes of large effect are likely to be found with low sample sizes, whereas decreasingly lower effect genes would only be found with larger and larger sample sizes. The rate of decline of α on number of accumulated genes was substantially lower for skeletal traits (*b* = –0.19 ± 0.05), *i.e*. height and waist related traits, than for the rest of traits (*b* = –0.48 ± 0.04) (see Figure 1 and Table S3[A]) probably because the former are less affected by selection.

**Fig 1:**
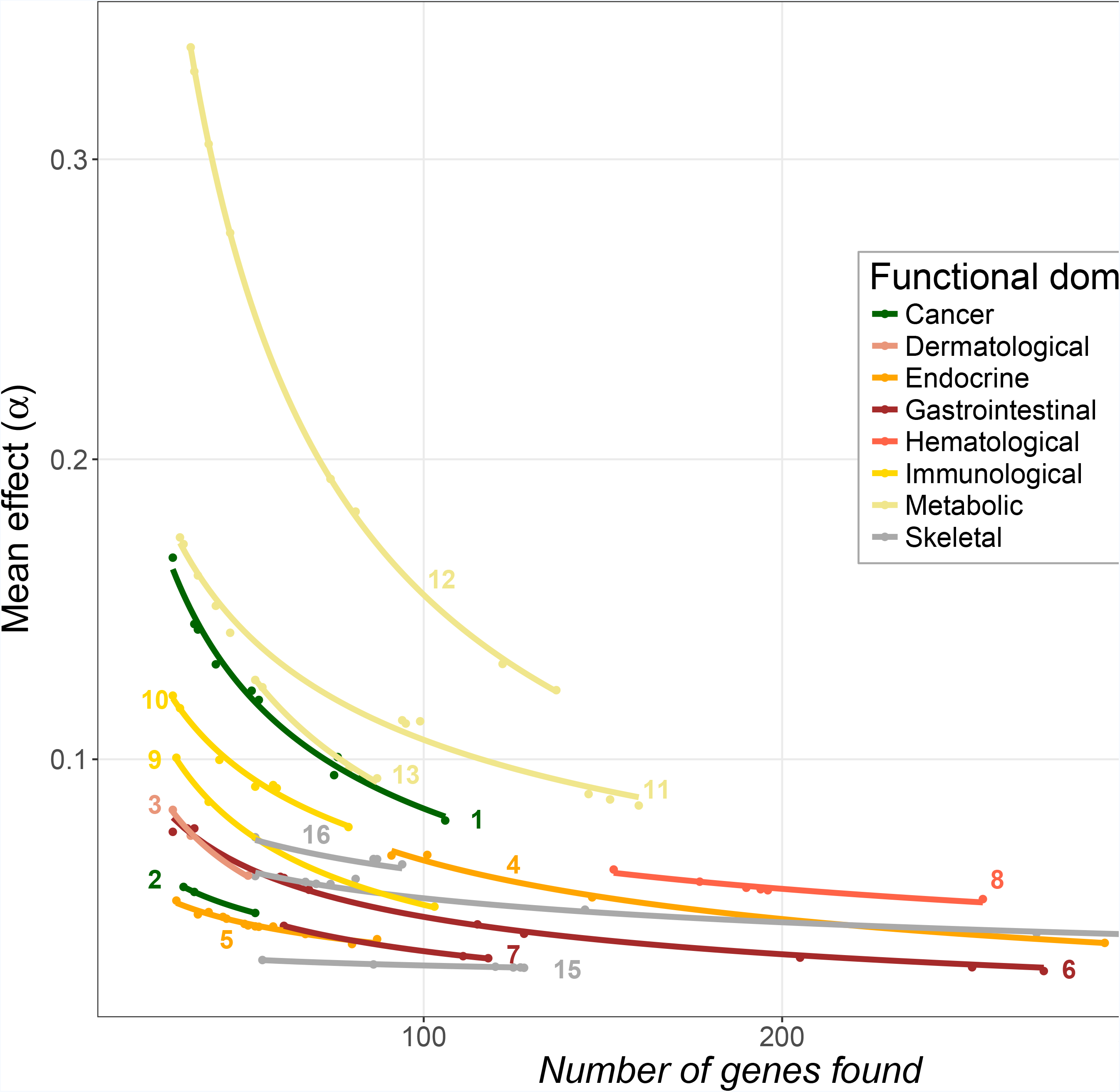
Decline of the average gene effect with the number of genes found. The points represents the cumulated results of successive GWA studies with increasing larger sample sizes. The first point at the left of the series is the mean effect of genes found in the GWAS with the lowest sample size (conditional on finding at least 30 genes), and the following give the mean effect of genes as additional ones are found by studies with higher sample sizes (usually, but not always, by more recent studies). The lines are the fit of the observations to an exponential model (average R2 = 0.96). Traits are colored depending on the functional domain they belong. Individual traits (i.e. lines) are: (1) Prostate cancer, (2) Testicular germ cell tumor, (3) Psoriasis, (4) Body mass index, (5) Type 2 diabetes, (6) Digestive disease, (7) Ulcerative colitis, (8) Neutrophil traits, (9) Rheumatoid arthritis, (10) Systemic lupus erythematosus, (11) Cholesterol, (12) HDL, (13) Triglycerides, (14) Height, (15) Waist-related traits, (16) Waist-to-hip-related traits.

Higher average gene effects were associated with lower allele frequencies for traits excluding skeletal ones (*b* = –0.49 ± 0.07), whereas this trend was non-significant for skeletal traits (*b* = 0.07 ± 0.07) (Table S3[D]). The change in average allele frequency with the number of genes found for each trait was generally positive (Table S3[E]), in accordance with the expectation that genes of small effect should be at higher frequencies to be detected. However, the average change was non-significant (*b* = 0.05 ± 0.05).

### Gene contributions to heritability

Estimates of the heritability explained by the additive contribution of individual genes 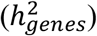 and of familiar heritability 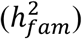 for 42 human traits are shown in Table S2, with averages 0.13 ± 0.02 and 0.50 ± 0.03, respectively. The proportion of 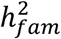 explained by 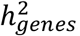 was 32% on average.

First, we wanted to know which genes contribute most to the heritability when these are classified by the magnitude of their effects. To do so, we established three arbitrary categories of gene effect sizes (low, medium and high) using 3-quantiles for each trait separately, where the same number of genes belongs to each quantile, and then measured its relative contribution to total 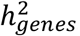. As a result, we found that for these 42 traits, on average, most of the heritability can be explained by genes of large effect (76.1%), with genes of medium and low effect explaining much lower proportions (16.7% and 7.2%, respectively) (Figure 2).

**Fig 2.**
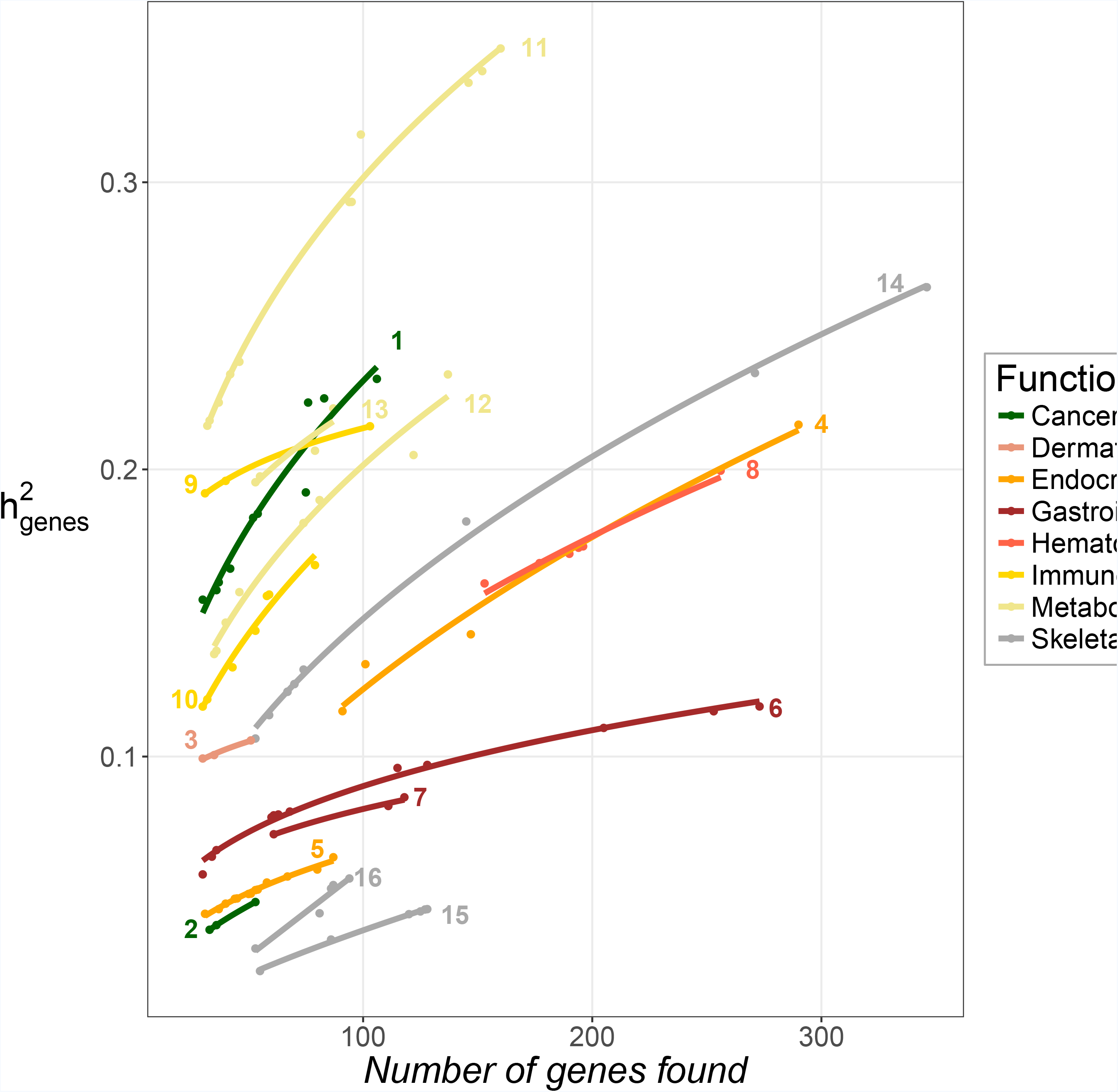
Relative gene contributions to heritability by effect size. Three arbitrary classes of gene effect sizes (high, medium and low effects) are assumed composed each of equal numbers of genes. Bars show the contribution of these classes to heritability: solid colors, low transparency and high transparency, for high, medium and low effect classes, respectively. Traits are ordered and colored by functional domain. Two dashed black lines highlight gene contributions of 33% and 67%, expected if each class would contribute proportionately the same to heritability.

Regression analysis in Figure 3 is shown again for a subset of 16 traits. It shows the explained heritability 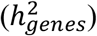 for each trait with an increasing number of genes found with increasing sample sizes (as for Fig. 1). A two-parameter exponential model gave the best fit to the data with average *R*^2^ = 0.97 (Table S3[F]). The figure shows that for most traits there is a substantial increase in the heritability explained as new genes have been found. However, for some traits (*e.g.* Digestive disease, number 6 in Figure 3) it looks like 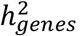 is approaching an asymptotic value. It can also be seen that in many cases the intercept is expected to be well above zero, suggesting that genes contributing most to the heritability were found in the studies with the lowest sample sizes, usually the earliest ones.

**Fig 3.**
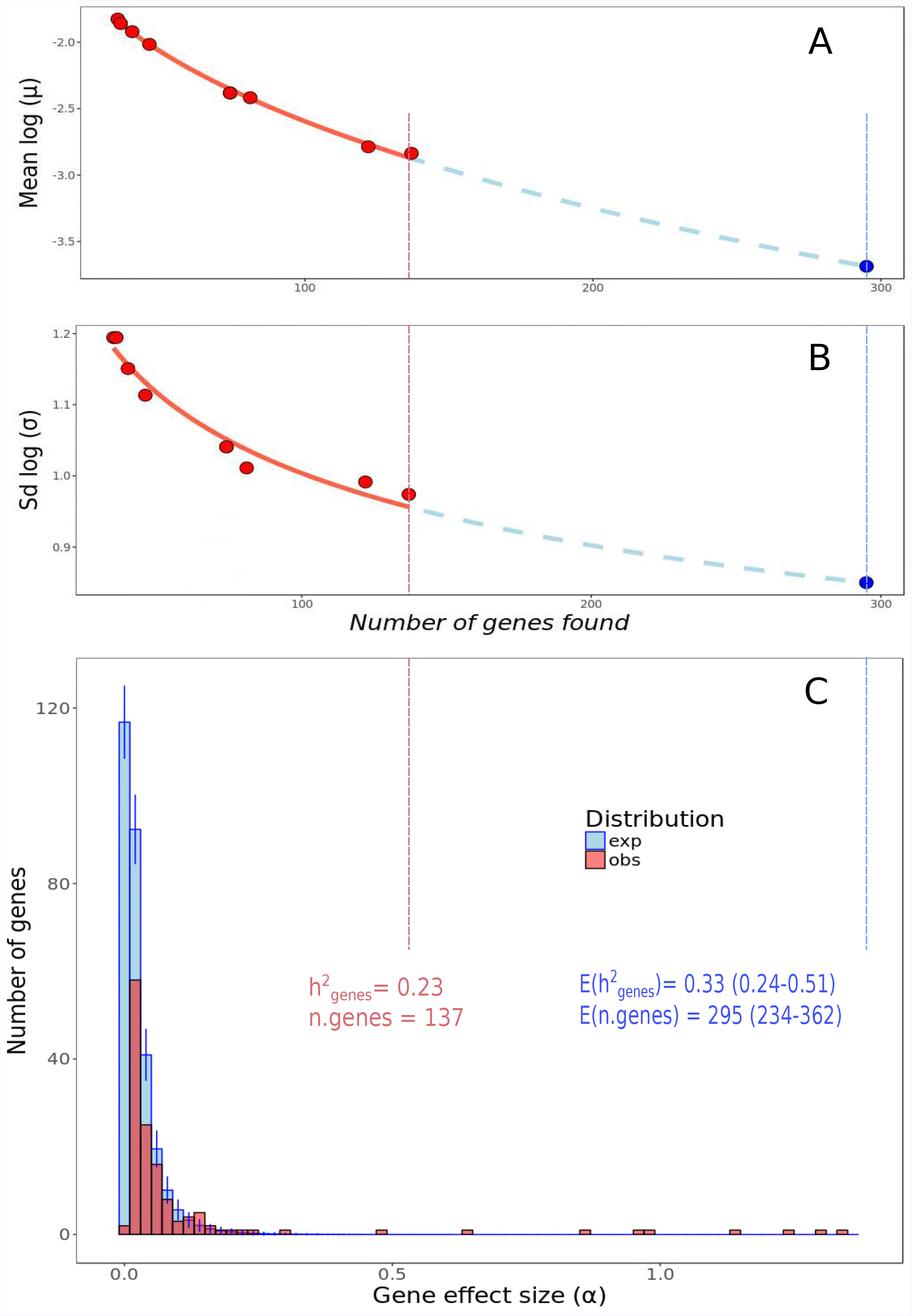
Increase of the heritability explained by genes found 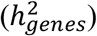 as the number of these is increasing. The points represent the observed values, while the lines are the fit to an exponential model (average R2 = 0.97). Traits are colored depending on the functional domain they belong. Individual traits (i.e. lines) are: (1) Prostate cancer, (2) Testicular germ cell tumor, (3) Psoriasis, (4) Body mass index, (5) Type 2 diabetes, (6) Digestive disease, (7) Ulcerative colitis, (8) Neutrophil traits, (9) Rheumatoid arthritis, (10) Systemic lupus erythematosus, (11) Cholesterol, (12) HDL, (13) Triglycerides, (14) Height, (15) Waist-related traits, (16) Waist-to-hip-related traits.

### Observed and expected distribution of effects, missing number of genes and asymptotic heritability

Gene effect sizes for most traits (90%) fitted better to a log-normal distribution than to any of the other distributions assessed (beta, exponential, gamma, gaussian, logistic, and uniform), the remaining 10% fitting best to a beta distribution (Table S4). Thus, we considered henceforth the log-normal distribution to make predictions of the expected distribution of gene effects.

Because the change of log-normal distribution parameters as new found genes are found fitted well to an exponential model (Figure 1 and Table S3[B,C]), we were able to predict them for any given number of genes (Figure S2 in Appendix 2), and thus infer the corresponding expected distribution of gene effect sizes. Using the relationship between gene effects and frequencies (Table S3[D]), we were further able to obtain the expected contributions to heritability. An illustration of this procedure is shown in Figure 4 for the trait HDL. Figs. 4A,B give the observed mean and standard deviation parameters of the log-normal distribution of effects (dots) for HDL genes as these are found and accumulated, and their expected projections under the exponential model fitted up to a number of genes which could explain the familiar heritability. Fig. 4C shows the observed distribution of effects with the current number of genes found by GWAS, and the corresponding expected distribution with a number of genes which would theoretically explain the familiar heritability for this trait. In this example, the heritability currently explained with 137 already found genes is 0.23, whereas the averaged familiar heritability reported in the Literature, 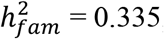, would be obtained with a range of 234-362 genes with a 95% confidence interval. This means that, if the familiar heritability of the trait HDL were explained by the additive contribution of single genes, between 97 and 225 new genes should be yet found. Note that the observed distribution shows a deficit of genes of small effect, but some genes of moderate effect are also missing.

**Fig 4.**
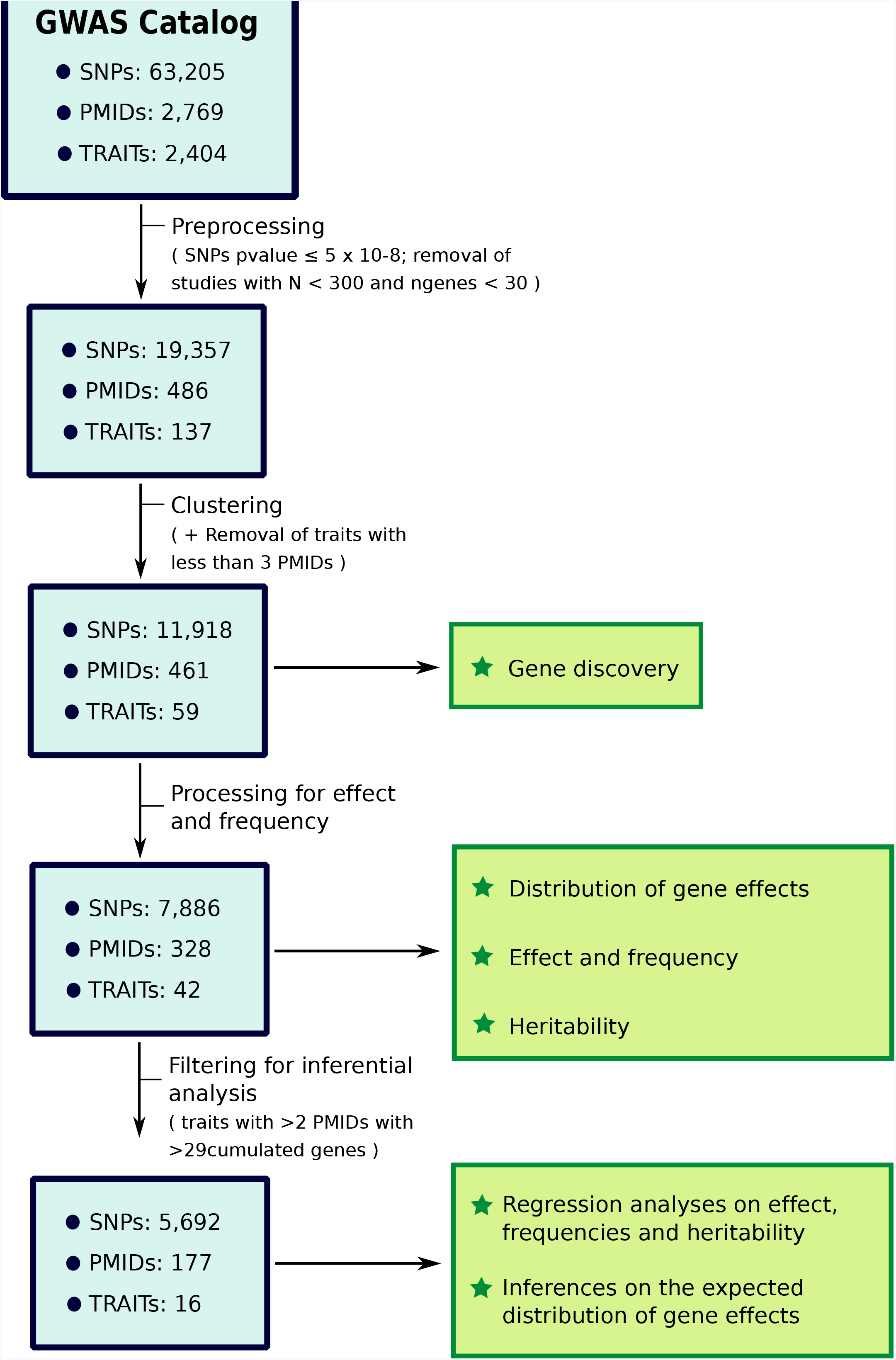
Illustration of the expected distribution inference process for trait HDL. Observed values are presented in red, and expectations in blue. Plots A and B show the change in the parameters of the log-normal distribution as new genes are being found: the mean parameter (A) and the standard deviation parameter (B). The blue point corresponds to the expected value of these parameters for the number of genes sampled from the expected distribution which gives the best approach to the familiar heritability. Plot C shows the distribution of gene effects, and the corresponding values of heritability and numbers of genes. Two vertical dashed lines connect observed (red) and expected (blue) distribution parameters with their corresponding values of heritability and number of genes.

We made this analysis for all 16 traits with sufficient information and the results are summarized in Figure 5, which shows the values of current values of heritabilities explained by the genes detected by GWAS (dark bars) and the familiar heritabilities (light bars). The heritability computed from the expected distribution of gene effects, which explains or approaches most to the familiar heritability, is shown as a dot (median value) and a 95% confidence interval. The expected distributions of effects obtained for all traits are shown in Figures S3 and S4, and the corresponding parameters in Table S6, as well as the expected number of genes (Appendix 2). For 11 out of 16 traits, the expected distribution is able to predict the familiar heritability accounting only for the contribution of additive effects of single genes. However, for the remaining 5 traits (body mass index, digestive disease, psoriasis, rheumatoid arthritis and type 2 diabetes), the familiar heritability cannot be reached when considering the additive effects of single genes. The expected distribution in these cases produces an asymptotic value of 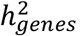 far below the average familiar estimate of heritability. The height of the error bar is highly related to the magnitude of the variance parameter (and therefore skewness) of the log-normal distribution. The number of genes required to reach either the familiar heritability or the asymptotic 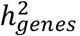 closest to the familiar one, is given on Fig. 5 legend, being on the order of hundreds or thousands.

**Fig 5.**
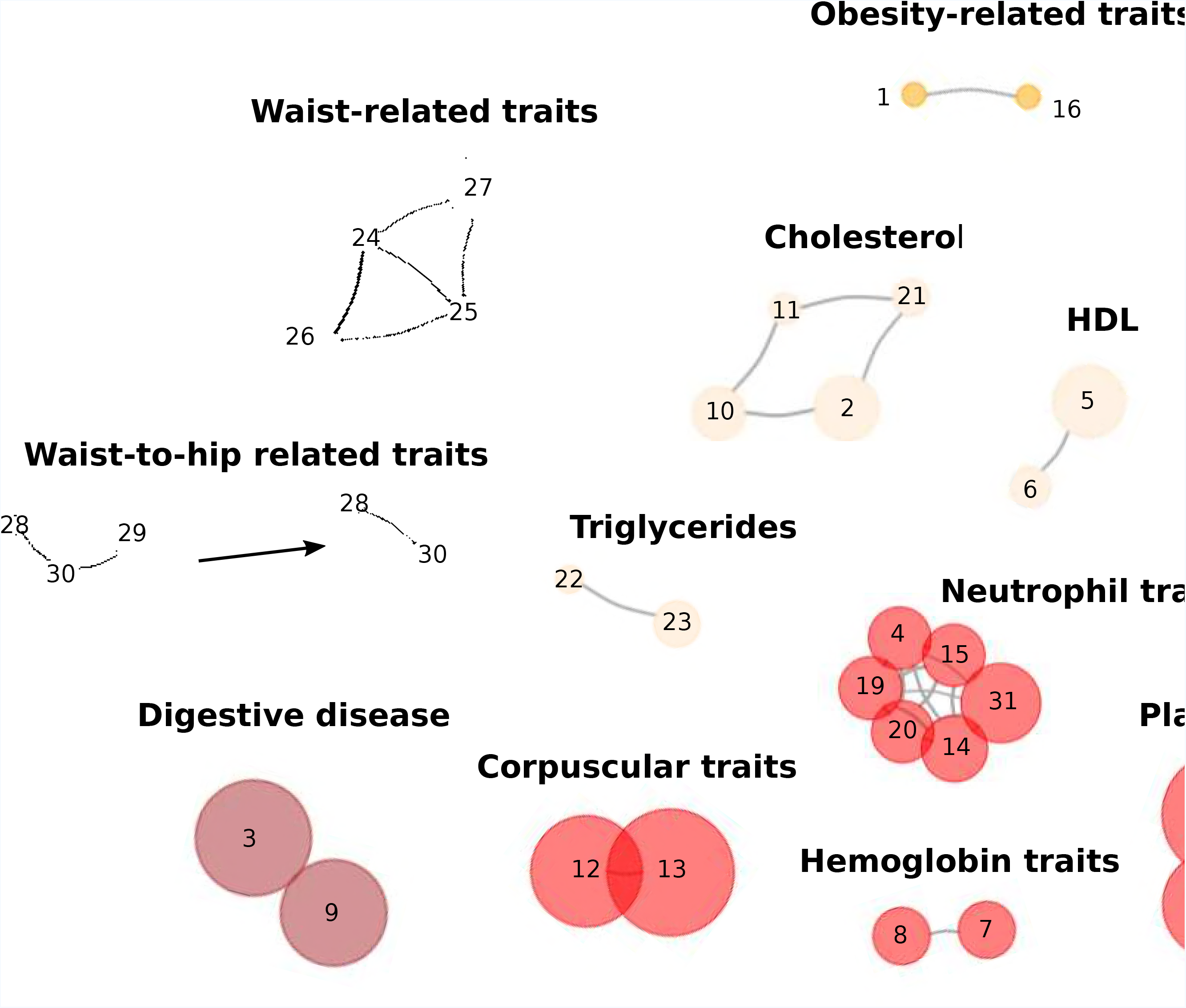
Observed and expected values of heritability. The full length of bars indicate the mean familiar heritability 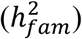 for the studied traits. In solid color it is shown the heritability explained by the variants already found and available from the Catalog 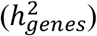. The blue error bar gives the inferred value of heritability (the dot corresponds to the median value) that approaches most to the familiar heritability with a 95% confidence interval, using data from the expected distribution of gene effects. The expected number of genes for each trait required to produce the expected heritabilities within the error bars are: Prostate cancer (106-256), Testicular germ cell tumor (1866-3705), Psoriasis (51-190), Body mass index (290-4287), Type 2 diabetes (547-1966), Digestive disease (273-842), Ulcerative colitis (118-1279), Neutrophil traits (256-290), Rheumatoid arthritis (103-265), Systemic lupus erythematosus (839-11789), Cholesterol (1433-4458), HDL (234-362), Triglycerides (87-530), Height (1138-1302), Waist-related traits (2047-2277), Waist-to-hip-related traits (247-377).

## DISCUSSION

By extracting the relevant data from the GWAS catalog we have been able to infer the main factors playing a role in the finding of risk variants for a number of diseases and common traits in humans. We have also been able to make predictions of the number and distribution of gene effects that could explain the missing heritability assuming the contribution of additive effects of single genes. For many of the studied traits this additive explanation is feasible whereas for some of them it can be rejected.

### Factors affecting new gene discovery

We first investigated the factors affecting new gene discovery. The factor with the strongest association (lowest P-value) to gene finding is population sample size, in agreement with the observation of an increase in statistical power detection for lower effect SNPs as sample size is increased(5,15). We found that, on average, about 53,000 individuals are required to start detecting genes, and about 6,000 for each new discovery, but with a wide range for the different traits (Table S5 in Appendix 1). From the data analysed it can be reckoned that a minimum sample size of about 120,000 individuals is, on average, enough to start finding variants with allele frequency lower than 0.1 and of small effect (defined as the third quantile of the distribution of effects; Fig. 2), a result coherent with other predictions(5).

Sample size and year of publication are, however, not the only relevant factors to be considered. We also analyzed the effect of using more than one population (or continent) of ancestry as a factor, and also genotyping platforms with a higher number of SNPs. While these factors are uncorrelated to population sample size, they have a significant effect on the number of new genes found. The size of the genotyping platform is an important determinant for new gene discovery as would be expected, with a significant regression of number of genes found on platform size. The number of continents sampled was also found to be a significant factor on gene discovery, and it is likely to be related to the discovery of ethnic-specific alleles. This emphasizes that human diversity in the form of different ethnics or populations of origin should be taken into consideration in GWAS designs. This is also required in order to accurately predict polygenic risk scores of diseases in different populations, as demonstrated by a recent study that showed that inferences made on non-European populations using European GWAS data were biased(16).

### Nature of the variation detected by GWAS

Our results indicate that the distribution of gene effects for human traits fits better to a log-normal distribution (Table S4) than to other common used distributions, including the gamma, widely assumed in population genetic studies(17–21). In the field of genetics, the log-normal distribution has been previously suggested by Drosophila DNA polymorphism data(22), and is usually assumed in models and natural processes arising not only in biology(23) but also in very different scientific disciplines(24,25).

We have also shown (Figure 1) that the average effect size tends to decrease for all traits studied as the number of discovered associated genes increases, supporting the idea that higher-effect mutations were discovered in the first GWAS (with lower sample sizes), while posterior analyses involving larger census sizes allowed discovering lower-effect mutations(11). It is worth mentioning that the slope of the decline in effect size with the number of genes was substantially lower for skeletal traits than for the other traits, generally associated or related to diseases. In addition, there was a significant negative relationship between the effect of genes and the allele frequency for non-skeletal traits, but this was non-significant for skeletal traits. This is in agreement with the hypothesis that genes of large effect associated to diseases are at lower frequencies because of natural selection. Even so, we found that there is a disproportionately larger contribution from genes of large effect to heritability, with genes of small effect contributing very little (Figure 2). This is in agreement with previous predictions(20) and contradicts models suggesting that most of the heritability for complex traits in humans must be due to genes of small effect(26).

### Expected distribution of gene effects and missing heritability

By fitting the change of the log-normal distribution parameters (mean and variance) for gene effect sizes, and the relationship between effects and frequencies, to an exponential model, we could infer the expected distribution of gene effect sizes that would be needed, if possible, to explain the familiar heritability for a restricted number of complex traits assuming an additive contribution of single genes. This also allowed us to infer the missing number of genes to reach the familiar heritability and their nature. Our results indicate that, for 11 out of 16 traits, the familiar heritability could be explained by the additive contribution of single genes (Figure 5). The number of genes required for this is below 500 for prostate cancer, ulcerative colitis, HDL, triglycerides, neutrophil traits and waist-to-hip-related traits, or about a few thousands for testicular germ cell tumor, systemic lupus erythematosus, cholesterol, height and waist-related traits. Thus, according to our results, a few hundred or a few thousand genes would be able to explain the missing heritability for this set of traits, assuming only additive contributions to heritability, in agreement with previous predictions(5).

For 5 out of 16 traits, however, also a few hundred or thousands of genes are required to reach an asymptotic value of heritability, but this is generally far below the familiar heritability (Figure 5). One anomalous result occurs with rheumatoid arthritis, for which the expected value of heritability was even below the currently observed value. The reason for this maladjustment is likely to be the bias generated when inferring the expected distribution. The observation that for some traits the expected heritability cannot reach the familiar one relies on the fact that, for these traits, the expected change in the shape of the distribution of gene effects predicts gene effect sizes too small to contribute significantly to heritability as the number of genes increase. In fact, the reason why some estimates of the missing number of genes to reach the familiar heritability are rather high in Figure 5 (*e.g.* for cholesterol, requiring 1433-4458 genes), is that the approach of the expected heritability to the familiar one is rather slow as the number of genes found increases because these have lower and lower effect sizes.

Our conclusion that for some traits the familiar heritability could not be retrieved by the accumulation of additive effects of single genes can also be deduced from Figure 3. If the increase in heritability with the accumulated number of genes is predicted from the figure for future numbers of genes to be found (regression parameters shown in Table S3[F]), it seems that for some traits, such as the five mentioned above (psoriasis, type 2 diabetes, digestive disease, body mass index, and rheumatoid arthritis) the heritability will reach a value far below the familiar one (Table S2).

We have made a number of assumptions which could affect the results obtained. First, we made a selection of the most informative SNPs available in the catalog for each trait, *i.e.* those with P-value ≤ 5 × 10^−8^. The reason was to consider only those for which the evidence of association with the trait is strong. This means that the number of genes assumed to be found is generally lower than that provided by the GWAS Catalog. With this assumption we would expect our predictions of number of genes and heritability from gene effects 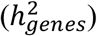 to be underestimations. Second, we took the SNP most significantly associated to a given gene (with the lowest P-value), and assumed that the estimated effect and frequency of that SNP is the same as for the corresponding associated gene. Thus, we assumed that the selected SNPs were at complete linkage disequilibrium with the causal gene. Therefore, the average effect size of the considered variants, and their contribution to heritability, would be expected to be overestimations. The estimated contribution to 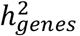 from the genes considered in the study are somewhat higher, but not very different, from those given in the Literature. In fact, the average 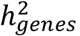 for all traits is, on average, a 32% of the familiar heritability (Table S2), a proportion of the order of those found in the Literature(27). This suggests that the two above sources of bias are not likely to be large, or they compensate to each other, giving support to our conclusions.

Our results show that hundreds to thousands of further loci are yet to be discovered for all traits studied. The expected distributions of effects inferred in this study (Figures 5, S3 and S4 in Appendix 2), show a main lack of genes of small effects to be found. This is also in agreement with the observation that the mean effect size monotonically decreases as more genes are being discovered (Fig. 1), in agreement with Park *et al*.(11). Therefore, it is expected that the missing heritability gap will be reduced very slowly with higher sample sizes and statistic power(27). However, not only variants of small effect are missing, and it is also expected to continue finding variants of moderate effect that have passed undetected so far, and that could still explain a part of the missing heritability.

It has been suggested that our inability to find the remaining loci in GWAS may come from a technical ground(9). Rare SNPs (say MAF < 5%) have a low coverage in current genotyping technology and are usually missing. Whole genome sequencing then could have the clue to find the proportion of missing heritability attributable to moderate or high effect loci, but it is expected that SNPs with extremely low frequencies contribute little to heritability, which has been already reported for diseases as Type 2 diabetes(28). In fact, simulation studies(20,29) have shown that full sequencing data accounting for SNP variation will not be able to increase substantially the estimates of heritability. However, it is possible that copy number variation, such as insertions and deletions that could be found by whole genome sequencing could make a substantial contribution to missing heritability(30–32). Furthermore, genome wide markers may overcome other statistical limitations for SNPs of complex traits, as inconsistent estimations of the gene effects due to SNPs in linkage disequilibrium with more than one QTL(33).

For traits for which an asymptotic heritability is expected to be substantially lower than the familiar heritability, other phenomena additional to the additive contribution of single genes may be involved. Epistasis has proven to be determinant in explaining heritability of several complex traits in yeast(34,35), and is becoming more evident that it might be a necessary element to explain heritability(36). In fact, epistasis has already been described playing an important role in psoriasis through the interaction of the HLA – ERAP1 loci(37) and other immunity disorders(38). Dominance could also be implied in heritability of many other traits, including height and BMI(14,39). Unfortunately, our results are limited to the additive component of heritability for single genes and cannot help further explaining its missing fraction beyond speculation.

Concluding, GWA analyses are a powerful tool to discover genes associated to complex diseases, and the success in finding the missing heritability may depend, in many instances, on our ability to detect low SNP effects with accuracy. For some traits, however, an additive contribution of single variants is not enough to explain the familiar heritability and other sources of variation must be sought.

## METHODS

All data manipulation, including statistical analysis, was carried out using the R language(40). The flowchart in Figure 6 summarizes the main processing and analytic steps. In short, we began by processing the GWAS Catalog in two steps, in order to get a set of data with the most meaningful information associated to SNPs and GWA studies. First, by filtering incomplete or low informative data, and creating new variables from the raw data in the Catalog, and second, by clustering together traits with a similar genetic background. This set was used for all analyses regarding the number of genes found by GWA studies. Additional analyses involving gene effect sizes, frequencies and contributions to heritability were performed on a more restricted set of data that required additional processing.

**Fig 6.**
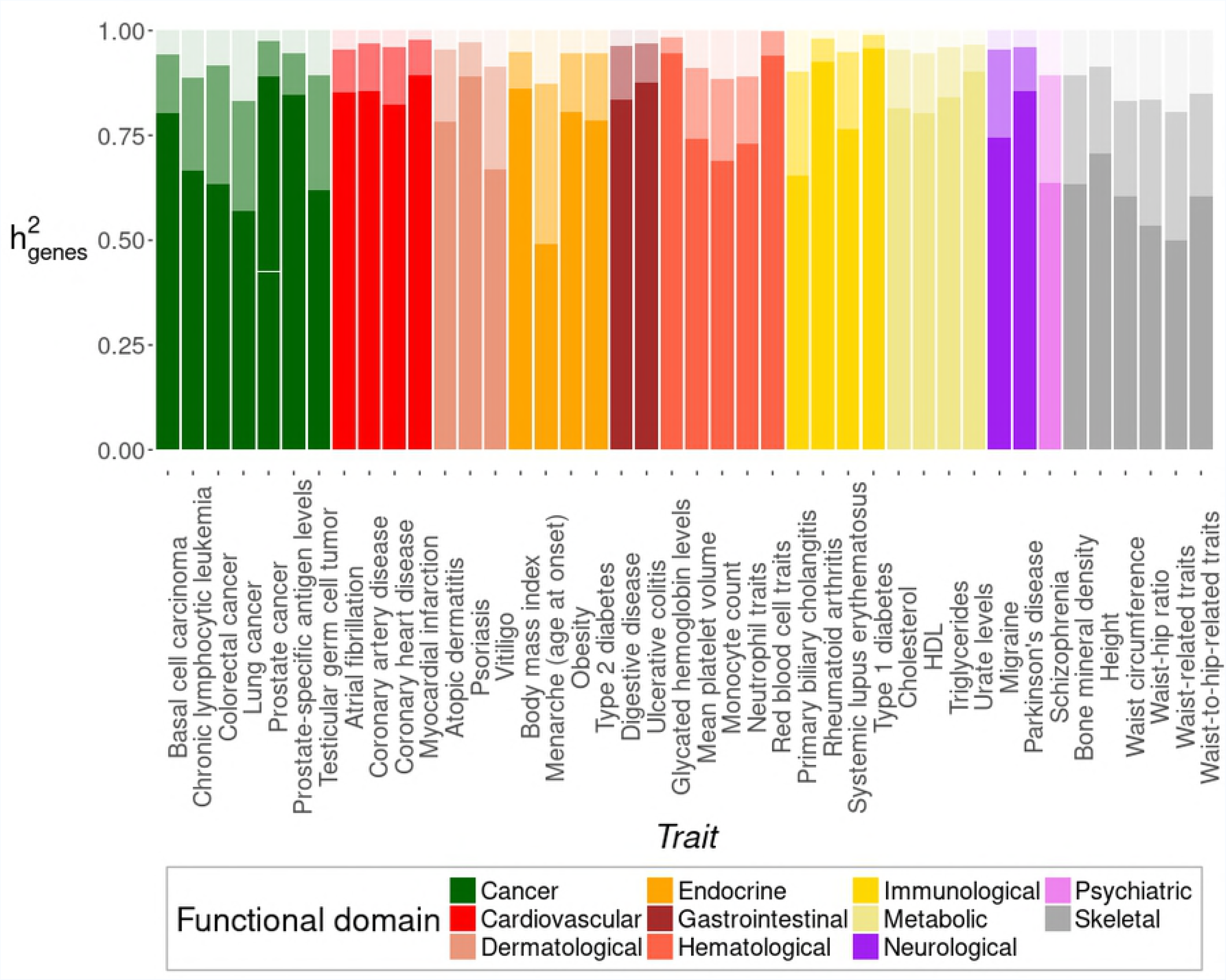
Flowchart summarizing the work schedule. The blue boxes refer to the main processing steps, indicating the number of SNPs, PMIDs and traits present in the dataset. Boxes in green highlight the analyses performed with the different datasets.

### Preprocessing of the GWAS Catalog

We worked with the NHGRI-EBI GWAS Catalog data(8) (publicly available at https://ebi.ac.uk/gwas/), accessed on the 05/12/2017, which consist on a database of results of published GWAS results that provides information on the SNPs detected and tens of fields including information on the mapped genes as well as the study design. The preprocessing of data is shown schematically in Figure 6. A glossary of all terms used are summarized in Table 1. We started by selecting a limited number of fields from the database for each scientific study PubMed ID (PMID), as the SNP ID itself, the mapped gene by the strongest SNP (*gene*), the effect, reported as an odd ratio or beta-coefficient (*effect*), the frequency of the risk allele (*q*) and the reported P-value (*pvalue*). PMID variables were also gathered, including the name of the disease or trait examined in the study (*trait*), the publication date (*date*), and information on the initial and replication samples, as well as information about the genotyping platform manufacturer used.

**Table 1:** Glossary of parameters and subscripts used in the Methods section.

|  |  |
| --- | --- |
| PMID | The PubMed identification number |
| trait | Diseases and traits examined in the study |
| SNP | Each single SNP reported in a given PMID |
| *gene* | The gene mapped to the strongest SNP. If the SNP is intergenic, it refers to the upstream and downstream genes |
| *N* | Total population sample size |
| *date* | Publication date |
| *year* | Publication year |
| *imputation* | A binary variable indicating SNP imputation |
| *nchip* | Number of SNPs passing QC |
| *ncont* | Number of continents of ancestry |
| *npops* | Number of populations of ancestry |
| *effect type* | The scale of the SNP effect sizes (BETA/OR) |
| *effect* | SNP effect size |
| BETA | Effect measured as beta-coefficient, in units of phenotypic standard deviations |
| OR | Effect measured as odd ratio |
| *pvalue* | The significance level (*i.e.* the reported P-value) |
| *q* | Risk allele frequency |
| $h_{SNP}^2$ | SNP-specific heritability |
| $h_{genes}^2$ | Heritability estimated with SNPs from the GWAS Catalog |
| $h_{fam}^2$ | Heritability estimates from family studies |

Additional variables were inferred from the above ones, as the year of publication (*year*) and others described below. First, the type of the effect (*effect type*) defined if the reported effect is an odd ratio (OR) or a beta-coefficient (BETA). The Catalog contains some ambiguity regarding the units of the effects registered. Doubtful cases were checked by looking at the corresponding publications and, if their effect could not be assigned as BETA or OR (*e.g*. because it was measured in trait units rather than in standardised units), it was marked as non-available (NA).

At the PMID level we could also infer, from the genotyping platform information, the number of SNPs passing QC (*nchip*) and if there was or not imputation of SNPs (*imputation*). Finally, we also calculated the total population sample (*N*) from the information of the initial and replication samples, as well as the number of populations (*npops*) and continents (*nconts*) with different ancestry considered.

We checked for the occurrence of a list of necessary variables (*effect*, *gene*, *pvalue*, *snp* and *trait*), and removed any row corresponding to a SNP without a complete information on these variables. We also limited our study to the most significant associations, eliminating SNPs with a significance level lower than the standard *pvalue* = 5 × 10^−8^. For all purposes only one SNP per gene (that with the lowest P-value) was considered. In addition, and because we wanted to investigate the distribution of gene effects with robustness, we only considered biological traits with at least 30 unique genes detected. After all these steps, the filtered catalog had information on a total of 19,357 SNPs observations, corresponding to 486 PMIDs and 137 traits (Figure 6).

### Clustering and filtering traits

We initially differentiated as many biological traits as unique names were given to the mapped disease or trait field in the original GWAS Catalog. However, it often occurs that different researchers studying the same trait publish their results using different trait names (*e.g*. “LDL” and “LDL levels”). In order to avoid working with duplicated or redundant traits, we clustered some of them on the basis of their common genetic background.

To do so, we first defined the following measure of similarity (*sim*):

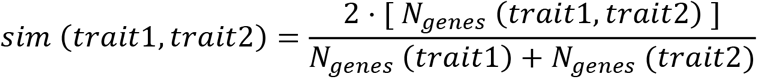

where *N*_genes_ refers to the number of unique genes found to be associated with a trait, or to two of them simultaneously. This measure ranges from 0 (no genes in common) to 1 (the same genetic background), as expected in the case where *trait1* equals *trait2*.

This let us build a similarity matrix with the pairwise similarities between traits. We only considered for clustering those traits sharing a high similarity (*sim* > 0.5), which represented the 98.85 quantile in similarity terms. This let us create graphs with traits acting as vertex, connected by edges between them always than *sim* > 0.5 (Figure 7).

**Fig 7.**
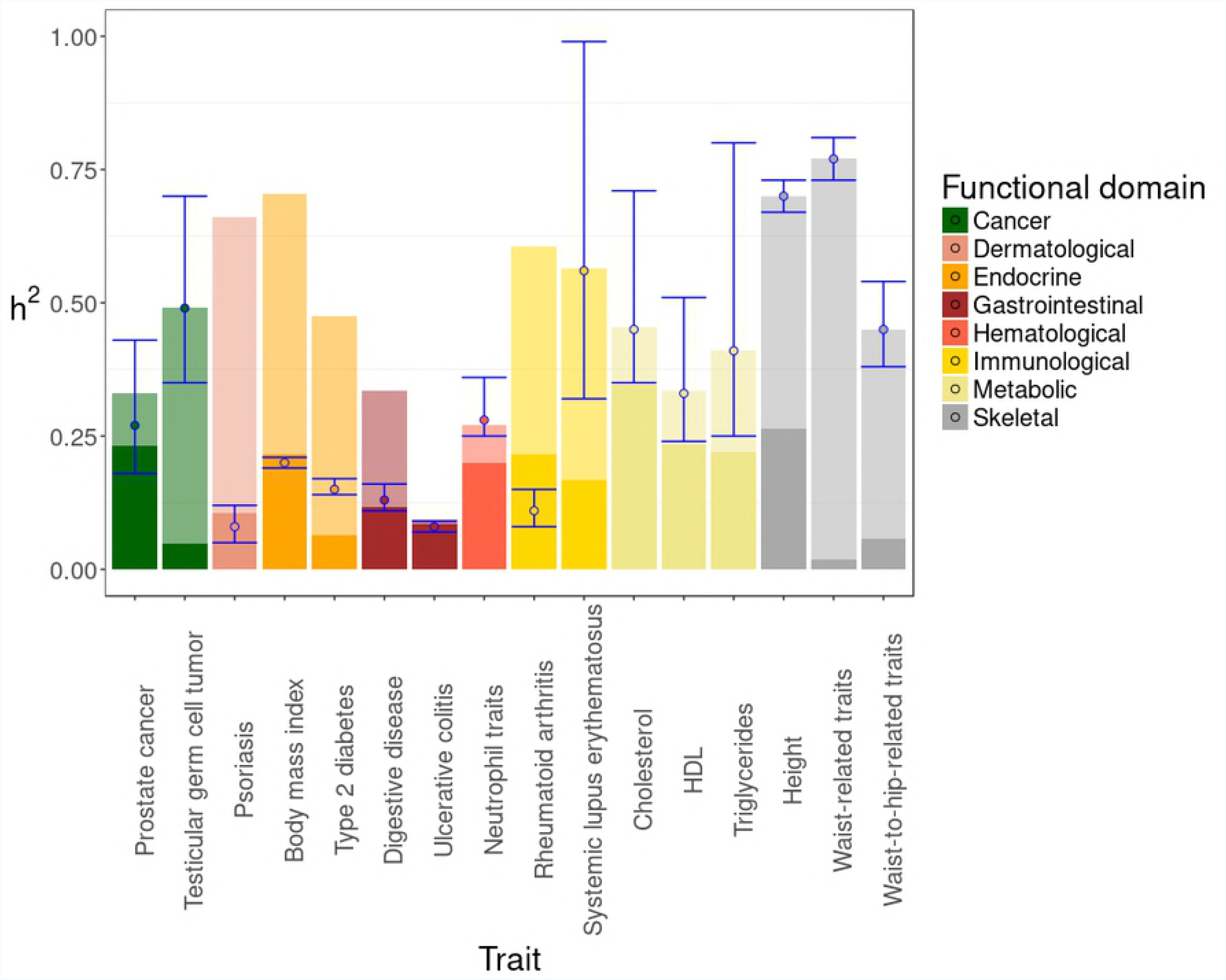
Disconnected graph showing traits (numbered vertex) that clustered together in groups (in bold text) with a high genetic similarity. The vertex color indicates the functional domain of the trait (orange: endocrine, khaki: metabolic, brown: gastrointestinal, red: hematological, grey: skeletal). The vertex size is proportional to the number of genes found for the trait. Individual traits are numbered as follows: 1 – Body mass index in physically active individuals; 2 – Cholesterol total; 3 – Crohn’s disease; 4 – Granulocyte count; 5 – HDL Cholesterol; 6 – HDL Cholesterol levels; 7 – Hematocrit; 8 – Hemoglobin concentration; 9 – Inflammatory bowel disease; 10 – LDL cholesterol; 11 – LDL cholesterol levels; 12 – Mean corpuscular hemoglobin; 13 – Mean corpuscular volume; 14 – Myeloid white cell count; 15 – Neutrophil count; 16 – Obesity; 17 – Platelet count; 18 – Plateletcrit; 19 – Sum basophil neutrophil counts; 20 – Sum neutrophil eosinophil counts; 21 – Total cholesterol levels; 22 – Triglyceride levels; 23 – Triglycerides; 24 – Waist circumference adjusted for BMI (adjusted for smoking behavior); 25 – Waist circumference adjusted for BMI in active individuals; 26 – Waist circumference adjusted for BMI in non-smokers; 27 – Waist circumference adjusted for body mass index; 28 – Waist-to-hip ratio adjusted for BMI (adjusted for smoking behavior); 29 – Waist-to-hip ratio adjusted for body mass index; 30 – White blood cell count.

In order to avoid spurious associations, we were very restrictive and only clustered together traits which belonged to graphs where every vertex was connected to more than 50% of the remaining vertex. When this condition was not satisfied, the less connected node (*i.e.* with less edges) was removed recursively until the above condition was met or the graph was completely disconnected. When we found two vertex with the same degree (*i.e*. number of edges), that with the lowest measure of average similarity was removed. This algorithm also allowed disconnected nodes to be reconnected if the above conditions were satisfied. For the purpose of analysis, the new clustered traits replaced the original ones. After this clustering step, traits studied by less than three different PMIDs were also removed. Table S1 shows all traits clustered, along with their *effect type*.

The database after processing and clustering included a total of 11,918 SNPs corresponding to 461 scientific studies for 59 human traits. It is worth mentioning that after the above preprocessing, clustering and filtering steps, the SNP/trait ratio and the PMID/trait ratio had been largely improved, the former from 26.3 to 202 SNP/trait and the later from 1.7 to 9.6 PMID/trait.

To simplify the representation of results, a second higher-level of clustering was carried out, with traits classified within functional domains similar to those used by Polderman *et al.*(41). The 14 functional domains considered were: Appearance, Cancer, Cardiovascular, Dermatological, Endocrine, Gastrointestinal, Hematological, Immunological, Metabolic, Neurological, Ophthalmological, Psychiatric, Respiratory and Skeletal. Table S1 also shows traits and clusters organized by these functional categories.

### Analysis of the factors associated to gene discovery

The analysis of the factors associated to gene discovery was carried out with the set of 59 traits obtained in the previous processing steps. We used ordinary least squares (OLS) multinomial linear regression in order to investigate the relationship between gene discovery and several factors of interest when planning a GWA study. These factors include information about the population sample, as its total size, *N* (initial and replication sample sizes), *npops* and *nconts*. The publication *year* of the study was also considered as a factor, along with other technical information such as *nchip* and *imputation*.

To avoid a possible bias because of the different genetic background of the traits considered, we performed the regression analyses over the residuals of a previous regression between the dependent variable and the traits(42,43). Thus, backward stepwise OLS multinomial linear regression was applied over these residuals up to removal of all non-significant factors. As some of the considered factors are likely to be not independent, we also estimated their partial correlation coefficients by the method of Pearson, using the ‘ppcor’ R package(44).

The most significant factor involved in gene discovery was total population sample size (*N*). We then analyzed, for each trait separately, the relationship between the accumulated number of genes discovered and population sample. The idea was that this regression analysis would inform on the population size required to discover new genes. The cumulative number of genes was calculated as follows. We sorted the scientific studies (PMIDs) by their population sizes. Thus, given the studies PMID1 and PMID2 for a given trait, with corresponding sample sizes *N*_1_ < *N*_2_, we identified the new genes found in PMID_1_, and then added to these the new genes found in PMID_2_ (*i.e.*, those which were not present in PMID_1_).

### Preprocessing for the analysis of gene effects and heritability

For the analysis of the effect sizes, risk allele frequencies and heritability, we removed from the database scientific studies where the *effect type* (BETA or OR) was ambiguous, corresponding to just six PMIDs. After revision of the publications, a total of 23 studies where the effects were measured in standard units rather than as beta coefficients were also identified and excluded. An additional study for type 2 diabetes contributing only one SNP with an extreme OR value, not reported in any other publication, was also excluded. Traits were pulled out from their clusters if they had different *effect type*. This was the case of the trait “Obesity”, which left the “Obesity-related traits” cluster shown in Fig. S2. SNPs without a measure of the risk allele frequency were also removed in this step.

Narrow-sense SNP-specific heritability 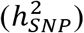 was estimated for the genes present in the GWAS Catalog through the calculation of their contribution to the additive variance 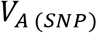 by using the classical formula(45):

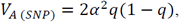

where *q* is the risk allele frequency of the most significant SNP associated to the corresponding gene, and *α* is the effect size of the SNP on the phenotype in standard deviation units.

For BETA traits, the additive variance equals the heritability 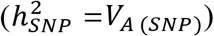, as the effects are measured in phenotypic standard deviations. For OR traits, we estimated the SNP-specific heritability 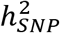 (*i.e*. variance in liability) following the method described by So *et al.*(46), assuming an additive effect of the SNP relative risk, and the prevalence values published in different epidemiology and genetic papers (see Table S2). From the 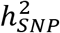 and frequency values, gene *effects* were obtained in the same units of phenotypic standard deviations as for BETA traits. Then, the total heritability estimated with SNPs from the GWAS Catalog 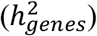 was calculated for each trait as the sum of all corresponding 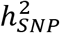 values.

After all the above filtering steps and checking for the minimum of 30 genes and 3 PMIDs per trait, the dataset for gene effects and heritability analyses had a total of 7,886 SNPs corresponding to 328 studies and 42 human traits (Figure 6). The estimated 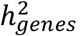 values are shown in Table S2 along with the reported values of familiar heritability 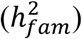 found in the Literature.

In order to measure the proportional contribution of different classes of gene effects to global heritability we defined three effect classes: low, medium, and high. These classes were assigned to each trait separately using 3-quantiles (*i.e*. terciles), so that the number of genes belonging to each effect class was the same.

### Analysis of the change in effect size, frequency, and explained heritability for increasing sample sizes

We assumed that gene effects and frequencies would be better estimated in studies with larger sample sizes. Thus, estimates obtained in studies with higher *N* were reassigned to the corresponding gene identity, independently of the study. That is, if we consider two studies, PMID_1_ and PMID_2_, regarding the same human trait, with census sizes *N*_1_ < *N*_2_, the gene effects and frequencies of genes found in PMID_1_ that were also present in PMID2 were assigned the values of the corresponding genes in PMID_2_.

We tested four different regression models to measure the relationship between variables in the analysis of gene effects, frequencies or heritability. These regression models were: simple linear regression: log_10_ *Y* = *a* + *b · X*; two-parameter exponential regression: *Y* = *a · X^b^*; three-parameter exponential regression: *Y* = *a* · *e^b·X^* +*c*; and four-parameter logistic regression 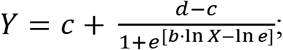 where the dependent variable *Y* may refer to the mean gene effect size, frequency, heritability or any other related variable such as the parameters of the distribution of gene effects, and the independent variable *X* is the number of genes found at a given stage in studies with increasing sample sizes.

When these models were performed using the accumulated number of genes as an independent variable, we only considered those traits that had at least 3 PMIDs in which the cumulated number of genes was at least 30, so that every measure point in the regression lines have at least 3 points, each corresponding to an estimate obtained with at least 30 genes. This corresponds to a subset of 16 traits, 177 PMIDs and 5,692 SNPs. For model selection, the Akaike Information Criterion (AIC) was used(47).

### Inferring the distribution of gene effects, the missing number of genes, and the asymptotic value of heritability

Gene effects were fitted into known probability distributions using the R package ‘fitdistrplus’(48) using the maximum likelihood estimation (MLE) method. In order to determine which distribution best fitted the observed gene effects, we considered the following possible continuous distributions: Beta, Exponential, Gamma, Gaussian, Logistic, Lognormal and Uniform. We then selected the best fit by using AIC. Adjustment to the fitted distribution was evaluated using the Kolmogorov-Smirnov test.

Given that the change in the parameters of the distribution of gene effects as new genes are being discovered could also be predicted with the regression parameters obtained in the previous analyses, the expected distribution of gene effects could be inferred. Thus, we obtained the distribution that allows explaining the observed value of familiar heritability 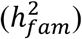, or the closest one, and thus the probability density of gene effects, including those yet unobserved. To do so, we assumed an increasing number of genes for each trait, and sampled that number of genes from the predicted distribution, with gene effect sizes and frequencies as expected from the regression parameters (Table S3). This process was repeated 10,000 times for each gene that was added, providing a distribution of expected heritability values. From this distribution, the parameters and numbers of genes which could explain 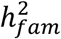 within 95% confidence intervals were chosen. If 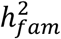 could not be explained by any expected distribution, the median heritability estimate closest to 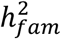 was chosen.

## ACKNOWLEDGMENTS

We are grateful to Enrique Santiago and Humberto Quesada for helpful comments. The analyses reported here were performed on the FinisTerrae machine provided by CESGA (Galicia Supercomputing Centre).

## SUPPORTING INFORMATION CAPTIONS

**Table S1.** Traits and diseases considered in this study, with their original name as seen in the GWAS Catalog, grouped by functional domain and cluster.

**Table S2.** Number of genes, prevalence (k), and heritability (h^2^) values for all traits and diseases considered in this study.

**Table S3.** Regression estimates related to gene effect sizes and frequencies, and heritability

**Table S4.** Parameters for the distribution of mutational effects for several human diseases and traits.

**Appendix1.** The factors determining new gene discovery. It includes **Figure S1** (Number of genes found for different studies depending on the sample size used) and **Table S5** (Regression estimates for the number of cumulated genes on the sample population size of the GWA studies).

**Appendix2.** The expected distribution of gene effects. It includes **Figure S2** (Change in the mean and standard deviation parameters of the log-normal distribution with the number of genes found), **Table S6** (Parameters of the observed and expected distribution of gene effects), **Figure S3** (Distribution of gene effects for non-disease traits) and **Figure S4** (Distribution of gene effects for disease traits).

